# Using dynamic biomaterials to study the temporal role of osteogenic growth peptide during osteogenesis

**DOI:** 10.1101/2023.07.19.549767

**Authors:** Fallon M. Fumasi, Tara MacCulloch, Julio Bernal-Chanchavac, Nicholas Stephanopoulos, Julianne L. Holloway

## Abstract

The extracellular matrix is a highly dynamic environment, and the precise temporal presentation of biochemical signals is critical for regulating cell behavior during development, healing, and disease progression. To mimic this behavior, we developed a modular DNA-based hydrogel platform to enable independent and reversible control over the immobilization of multiple biomolecules during in vitro cell culture. We combined reversible DNA handles with a norbornene-modified hyaluronic acid hydrogel to orthogonally add and remove multiple biomolecule-DNA conjugates at user-defined timepoints. We demonstrated that the persistent presentation of the cell adhesion peptide RGD was required to maintain cell spreading on hyaluronic acid hydrogels. Further, we discovered the delayed presentation of osteogenic growth peptide (OGP) increased alkaline phosphatase activity compared to other temporal variations. This finding is critically important when considering the design of OGP delivery approaches for bone repair. More broadly, this platform provides a unique approach to tease apart the temporal role of multiple biomolecules during development, regeneration, and disease progression.

## Introduction

The extracellular matrix (ECM) provides key structural and biochemical signaling cues to nearby cells. The ECM is highly dynamic, and the precise presentation of these signaling cues in space and time is responsible for regulating cell fate during development, homeostasis, wound healing, and disease progression.^1–3^ The dynamic nature of the ECM is of particular interest in the context of wound healing, where the ability to mimic this behavior using tissue-engineered scaffolds is critical for functional tissue regeneration.^4–6^ As an example, bone has four temporal stages of fracture repair, with three main groups of biochemical signaling cues – inflammatory cytokines, morphogens, and angiogenic factors – that play a key role in regulating these stages.^7,8^ The presentation and expression level of these signaling cues dynamically changes as healing progresses from one stage to the next stage. However, it is an ongoing challenge to replicate the vast number and temporal complexity of these signaling cues using biomaterial-based approaches.

The importance of recapitulating the dynamic nature of the ECM has resulted in a suite of biomaterials that can be spatiotemporally controlled with either molecular or physical triggers.^9–12^ A range of materials have demonstrated the ability to modulate ligand presentation^13–15^ and hydrogel stiffness.^16–18^ Nonetheless, few of these approaches allow for the orthogonal and completely reversible addition and removal of multiple (three or more) biochemical signals, which is needed to fully recapitulate the dynamic complexity of the ECM. For example, light is one of the most powerful triggers for changing material properties—usually through photocleavage, or photo-triggered chemical reactions—due to its exquisite spatial and temporal control. However, despite elegant work controlling different types of signals (e.g., peptide or protein presentation) with light,^15,17,19^ the chemical moieties necessary for such orthogonality can be challenging to synthesize, and spectral overlap generally limits the maximum number of signals that can be tuned to three. Furthermore, while signal removal is reasonably straightforward using photocleavage reactions, reintroducing multiple signals with different wavelengths of light is still challenging.

To better mimic the dynamic nature of the ECM, it is necessary to design biomaterials that can reversibly and orthogonally control multiple biochemical signaling cues. DNA hybridization has shown great promise as a tool for the reversible presentation of peptides and proteins^20^ or reversible tuning of hydrogel stiffness by breaking and reforming crosslinks.^21^ The advantage of DNA as a tether is that hybridization can be reversed by introducing a single-stranded DNA “toehold”;^22,23^ where addition of a fully complementary strand will hybridize to the toehold and, via branch migration, displace the original hybridized strand. In this way, multiple signals can be controlled independently by using different sets of orthogonal DNA strands.

Here, we describe the use of reversible DNA handles to dynamically control bioactive peptides (Figure 1A). Orthogonal sets of complementary oligonucleotides were designed to selectively add and remove multiple biomolecule signals over several cycles. The initial DNA strand was covalently tethered onto a hyaluronic acid (HA) hydrogel using thiol-norbornene photochemistry, which enables spatial control over the conjugation reaction via photopatterning. HA is a key component of the extracellular matrix, interacts with cell receptors CD44 and RHAMM, and can easily be modified to incorporate chemical functional groups of interest (i.e., reactive groups for crosslinking or biomolecule immobilization).^24,25^ Additionally, HA does not innately support integrin-mediated cell adhesion, which enables us to precisely control cell-matrix adhesion through the synthetic incorporation of cell-adhesive peptides. The complementary peptide-DNA conjugate was added to the hydrogel and immobilized via hybridization. The peptide-DNA conjugate was removed via addition of the complementary displacement DNA via toehold mediated displacement. The specificity and reversibility of this platform was demonstrated using fluorophore labeled DNA. To demonstrate the utility of this platform, we evaluated the temporal in vitro cell response to two peptides of interest: the cell-adhesive peptide, RGD, and osteogenic growth peptide (OGP). RGD is derived from fibronectin and is commonly used to promote cell adhesion onto biomaterial scaffolds via integrin binding.^26–28^ In vitro cell morphology was measured as a function of RGD concentration and immobilization time. OGP is a naturally occurring growth factor and is known to enhance proliferation and osteogenic differentiation.^29–31^ The temporal role of OGP during in vitro osteogenesis was evaluated using an alkaline phosphatase (ALP) assay. Both studies demonstrated the importance of biomolecule temporal presentation on modulating in vitro cell behavior. Thus, this platform provides a unique approach to tease apart the temporal role of multiple biomolecules during development, regeneration, and disease progression.

**Figure 1:**
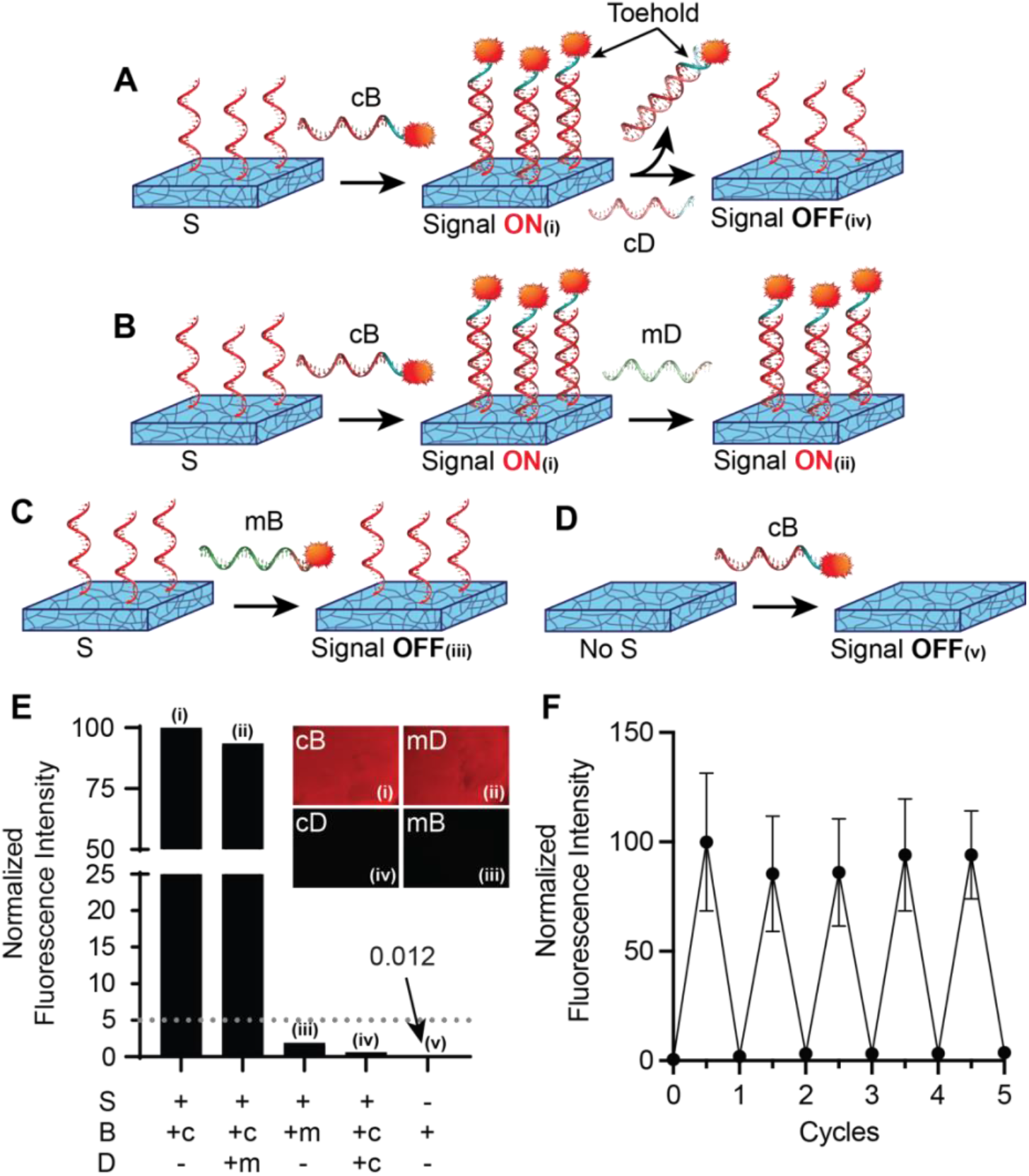
Reversible DNA handles enable highly specific biomolecule addition and removal at user-defined timepoints over multiple cycles. (A) First, surface (S)-DNA was photo-conjugated to the hydrogel. At user-defined times, the complementary biomolecule (cB)-DNA was added to turn the signal ON and the complementary displacement (cD)-DNA was added to turn the signal OFF via toehold mediated displacement. This approach was highly specific: (B) mismatched displacement (mD)-DNA was unable to remove cB-DNA, (C) mismatched biomolecule (mB)-DNA was unable to hybridize to S-DNA, and (D) cB-DNA did not bind to the hydrogel without S-DNA. (E) cB/mB-DNA was labeled with rhodamine and fluorescence intensity was measured at various stages of addition and removal, as indicated by the lower case Roman numerals within the schematic. (F) Fluorescence intensity was measured as cB-DNA and cD-DNA were added to the hydrogel to turn the signal ON and OFF, respectively, and repeated for five cycles to demonstrate reversibility (n=4). Error bars represent standard deviations from the mean.

## Materials and Methods

### Materials

All chemicals were purchased from Sigma-Aldrich unless otherwise noted.

### Norbornene-Functionalized Hyaluronic Acid (NorHA) Synthesis

A two-step protocol was performed to synthesize norbornene-functionalized hyaluronic acid (NorHA).^32,33^ First, the tetrabutylammonium salt of hyaluronic acid (HA-TBA) was synthesized by exchanging the sodium salt of hyaluronic acid (HA, Lifecore Biomedical, 60 kDa) using the Dowex 50W×4 (50-100 mesh) ion exchange resin followed by neutralization with 0.2 M tetrabutylammonium hydroxide (TBA-OH). HA-TBA was frozen, lyophilized, and analyzed with ^1^H NMR. Secondly, HA-TBA was combined with 5-norbornene-2-methylamine (Tokyo Chemical Industry) and benzotriazole-1-yl-oxy-tris-(dimethylamino)-phosphonium hexafluorophosphate (BOP), dissolved in anhydrous dimethyl sulfoxide (DMSO), and the reaction vessel was purged with nitrogen gas. The solution reacted with stirring for 2 hours at room temperature to form NorHA. Following reaction, NorHA was purified via extensive dialysis using deionized (DI) water at room temperature, frozen, lyophilized, and analyzed using ^1^H NMR to characterize norbornene functionalization (Figure S1). NorHA was synthesized to yield a norbornene functionalization of approximately 65%.

### NorHA Hydrogel Formation

Glass slides were modified with thiol-containing silicone (SYLGARD 184, Dow) to bind the hydrogel to the surface and provide a stable surface for easy handling. SYLGARD components were mixed according to manufacturer instructions and 10 uL of (3-mercaptopropyl)trimethoxysilane (MPTS) per 1 g SYLGARD was added to the mixture. Glass slides were coated in a thin layer of the SYLGARD/MPTS mixture and reacted for 1 hour at 125 °C. Modified slides were washed sequentially in dichloromethane, ethanol (Fisher Scientific), and DI water; dried; and stored under vacuum until used. To form hydrogels, 4% w/v NorHA was dissolved in phosphate-buffered saline (PBS, Fisher Scientific) containing 0.05% w/v photoinitiator (Irgacure 2959, I2959) and the non-degradable crosslinker 1,4-dithiothreitol (DTT). DTT was added at a stoichiometric molar ratio of 0.2:1 thiol to norbornene functional groups, leaving 80% of the original norbornene groups available for photoconjugation at a later step.

The pre-gel solution was transferred into a custom silicone mold (SYLGARD 184, Dow), covered with a thiolated glass slide (thiol side down), and crosslinked using ultraviolet (UV) light at 10 mW/cm^2^ (320-390 nm, OmniCure S1500) for 5 minutes. After crosslinking, the silicone mold was removed, and the hydrogels remained attached to thiolated glass slides.

### Peptide Synthesis

All peptides were synthesized using standard fluoren-9-yl methoxy carbonyl (Fmoc) solid-phase peptide synthesis. RGD was synthesized on a rink amide AM resin (100-200 mesh, 0.57 mmol/g) and OGP was synthesized on a pre-loaded glycine wang resin (100-200 mesh, 0.42 mmol/g). Peptides were cleaved from the resin using a 90:5:2.5:2.5 mixture of trifluoroacetic acid (TFA):2,2’-(ethylenedioxy) diethanethiol (DODT):triisopropyl silane (TIPS):water and precipitated using cold diethyl ether. The peptides were purified using reverse phase high-performance liquid chromatography (HPLC) (Waters, Phenomenex C18 column) with a water/acetonitrile gradient (0-80% acetonitrile over 60 minutes) and 0.1% TFA. Purified peptides were lyophilized, stored at -20 °C, and analyzed using matrix-assisted laser desorption/ionization time-of-flight (MALDI-TOF) mass spectrometry (Bruker Microflex LRF). The following peptides were synthesized: ZGYG<u>RGDSP</u>G (RGD, bioactive domain is underlined)^33,34^ and ZGGCG<u>ALKRQGRTLYGFGG</u> (OGP, native sequence is underlined).^29,35^ Z represents the unnatural amino acid azidolysine, which was used for conjugation to DNA.

### Peptide-DNA Conjugate Synthesis

All peptide-DNA conjugates (Table 1) were synthesized via the reaction scheme in Supplementary Figure S2A and as previously described.^36^ Briefly, oligonucleotides with a terminal amine moiety (Integrated DNA Technologies) were dissolved to 500 µM in 10 mM PBS (pH 7.5), 20 molar equivalents of dibenzocyclooctyne-sulfo-*N*-hydroxysuccinimidyl ester (DBCO-sulfo-NHS ester) was added, and the resulting solution was allowed to react for 3 hours at room temperature. The DBCO-modified DNA was purified using reverse phase HPLC (Agilent 1220 Infinity, Zorbax Eclipse XDB C18 column) with a linear gradient of 50 mM triethylammonium acetate/methanol (0-80% methanol over 45 minutes). Absorbance peaks at 260 and 309 nm were monitored, and peaks over a 200 mAU threshold were collected and concentrated using a 3 kDa MWCO spin filter. The fraction that contained both absorbance peaks, 260 nm for the DNA and 309 nm for the DBCO, was mixed with the desired azidolysine-containing peptide in 10 mM PBS at a 1:4 molar ratio of DNA:peptide, and allowed to react overnight at room temperature. The peptide-DNA conjugate was purified using the same HPLC method as the DBCO-modified DNA, the buffer was exchanged with PBS, and stored frozen at - 20 °C until further use. Peptide-DNA conjugates were analyzed using MALDI-TOF mass spectrometry with 6-aza-2-thiothymine as a matrix. Four orthogonal sets of DNA sequences were used as indicated in Table 1, where each set included a surface (S)-DNA, complementary biomolecule (cB)-DNA, and complementary displacement (cD)-DNA strand. Mismatched biomolecule (mB)- and mismatched displacement (mD)-DNA strands were also synthesized to serve as controls and evaluate hybridization specificity.

**Table 1:**
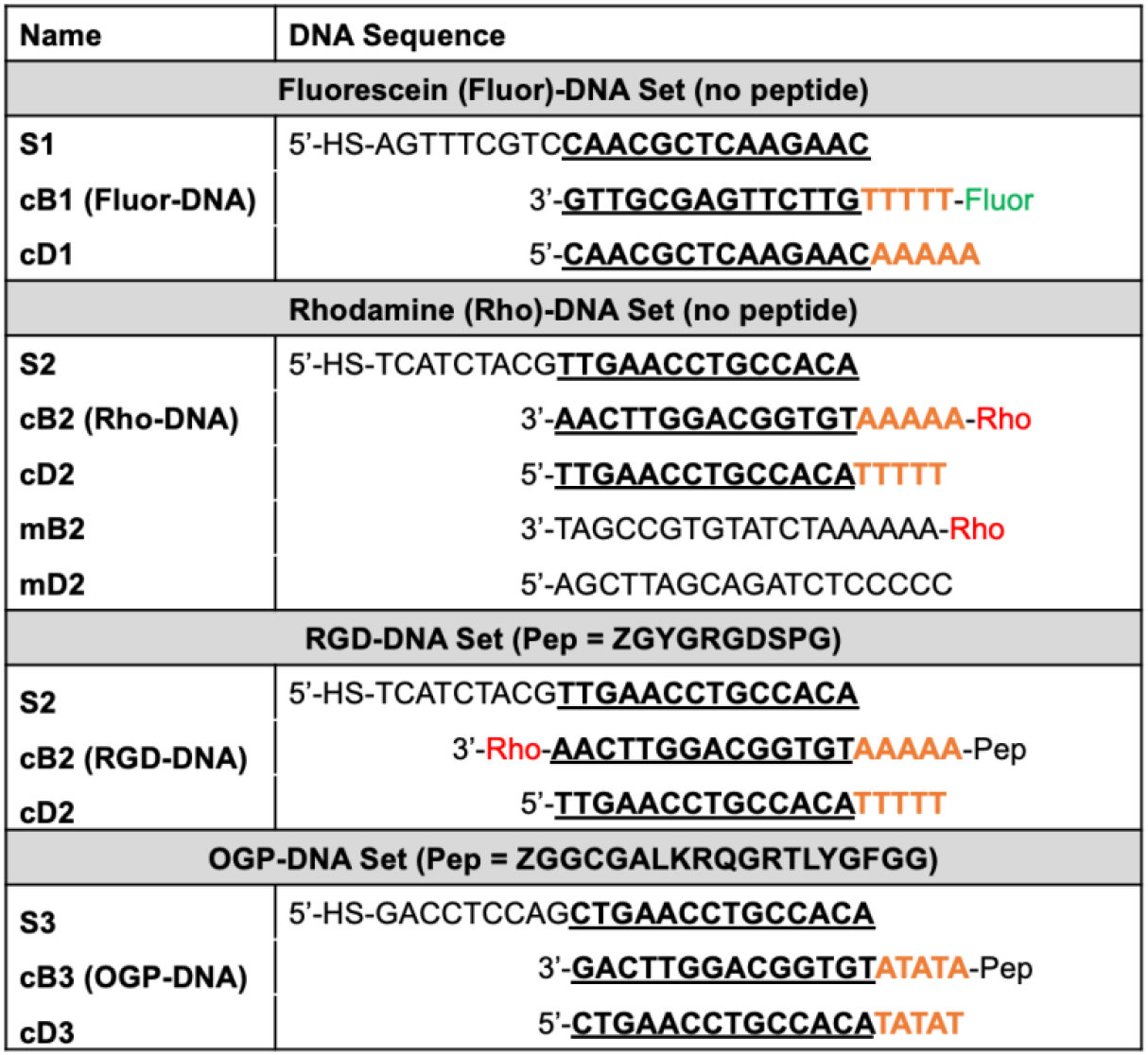
Nucleotide sequences for all DNA strands used. The complementary nucleotide sequence between S-, cB-, and cD-DNA is bold and underlined. The complementary toehold region between the cB- and cD-DNA is bold and orange. Abbreviations: surface (S), complementary biomolecule (cB), complementary displacement (cD), mismatched biomolecule (mB), and mismatched displacement (mD).

### Addition and Removal of Peptide-DNA Conjugates on NorHA Hydrogels

NorHA hydrogels were crosslinked with a 0.2:1 thiol to norbornene molar ratio as described in the NorHA Hydrogel Formation section. A solution of the surface (S)-DNA (50 μM of S1 or S2 in PBS) containing 0.05% w/v I2959 was added dropwise to the hydrogel surface, allowed to soak for 5 minutes, and exposed to UV light for 1 minute at 10 mW/cm^2^ to covalently photo-conjugate S-DNA to NorHA. Unconjugated S-DNA was removed via washing with PBS. A solution of the complementary biomolecule (cB)-DNA (50 μM of cB1 or cB2) was added dropwise to the hydrogel surface and soaked for 5 minutes to allow hybridization to S-DNA. Unbound cB-DNA was removed via washing with PBS. Bound cB-DNA was visualized using fluorescent microscopy (Leica DMI6000 B). The cB-DNA was engineered to contain a bioactive peptide, a toehold region for displacement, and a fluorophore for imaging. A solution of the complementary displacement (cD)-DNA (100 μM of cD1 or cD2) was added dropwise to the hydrogel surface and soaked for 30 minutes to enable cB-DNA removal via toehold-mediated strand displacement. Following removal, hydrogels were washed with PBS and cB-DNA removal was confirmed using fluorescent microscopy. Experimental groups with mismatched biomolecule (mB)- and mismatched displacement (mD)-DNA stands, as well as without S-DNA on the surface were used as controls to demonstrate the specificity of this hybridization scheme.

### Thiol Quantification Assay

The efficiency of the norbornene-thiol reaction was measured using a thiol quantification assay (Measure-iT Thiol Assay Kit, Invitrogen).^37^ The cell adhesive, RGD peptide (containing a cysteine amino acid residue for attachment) was used as a model thiol-containing molecule for this study. RGD was dissolved in PBS containing 0.05% w/v I2959 to create 25, 50, 100, or 500 µM RGD solutions (n=3). The RGD solution was added dropwise to the hydrogel surface and immediately exposed to UV light for 60 seconds at 10 mW/cm^2^ to attach RGD to NorHA. The solution was removed from the hydrogel surface and the remaining thiols were quantified using a standards curve and the thiol assay kit according to the manufacturer’s protocol. The tethered RGD concentration was calculated by subtracting the remaining thiols in the RGD solution from the initial RGD solution concentration.

### Compressive Testing

A bench top mechanical testing system (Instron Materials Testing System Series 5943, 50 N load cell) was used to test hydrogels in unconfined compression at room temperature.

Cylindrical samples approximately 4.8 mm in diameter and 4.3 mm in height were compressed at 20% strain per minute. The compressive modulus was calculated from the initial linear region of the stress vs strain curve. Three hydrogel experimental groups were evaluated: hydrogel with no DNA; hydrogel containing photo-conjugated S1-DNA and S2-DNA; and hydrogel containing photo-conjugated S1-DNA and S2-DNA and hybridized with cB1-DNA and cB2-DNA, respectively (n=6).

### In Vitro Cell Culture

Human mesenchymal stem cells (hMSCs) (Lonza PT-2501) at passage 3 were used for all in vitro cell culture studies. Cells were cultured in growth media (α-Minimum Essential Medium (MEM) without nucleosides (Gibco) supplemented with 16% fetal bovine serum (FBS, Gibco), 1% L-glutamine (Gibco), and 1% penicillin-streptomycin (pen/strep, Gibco)) at 37 °C and 5% CO_2_. Prior to cell seeding, all hydrogels were sterilized via ethanol immersion for 30 minutes followed by extensive washing using sterile PBS. Additionally, all hydrogels contained 10 μM RGD peptide in the bulk phase, which was added to the pre-gel solution and tethered to NorHA during the hydrogel crosslinking step. A very low level of RGD was added to all hydrogel groups to ensure cell attachment while minimizing cell spreading, as HA does not innately support integrin-mediated cell adhesion. RGD concentrations mentioned in the next section are in addition to the 10 μM RGD peptide in the bulk phase.

### Cell Morphology

For cell morphology studies, cells were seeded onto the hydrogels at 5,000 cells/cm^2^ and cultured in growth media for up to 7 days. Two sets of experiments were performed: persistent RGD immobilization and dynamic RGD immobilization. For persistent RGD immobilization, cells were seeded onto hydrogels containing no RGD (negative control), 100 μM RGD (no DNA control), 100 μM RGD-DNA + 100 μM S-DNA, or 1000 μM RGD (positive control). After 3 days, samples were fixed, and cell morphology was analyzed (n=3). For dynamic RGD immobilization, cells were seeded onto hydrogels containing 100 μM RGD-DNA + 100 μM S-DNA. On day 3, RGD-DNA was allowed to remain or removed via the addition of 200 μM cD-DNA. After 7 days, samples were fixed, and cell morphology was analyzed (n=3). For all groups, the RGD peptide alone or S-DNA was photo-conjugated to the hydrogel using the previously described method. The complementary RGD-DNA was added to the hydrogel following photo-conjugation of S-DNA. Peptide immobilization was performed after sterilization and prior to cell seeding.

Cell morphology was evaluated using rhodamine-phalloidin (Invitrogen) and 4’,6-diamidino-2-phenylindole dihydrochloride (DAPI, Invitrogen) staining to visualize the cytoskeleton and nucleus, respectively. Samples were imaged using an Olympus BX63 microscope and ImageJ was used to quantify cell number and area.

### Alkaline Phosphatase Activity

For osteogenesis studies, cells were seeded onto the hydrogels at 25,000 cells/cm^2^ and cultured in growth media for 24 hours. After 24 hours, the growth media was exchanged with osteogenic induction media (osteomedia, StemXVivo, Bio-Techne) (Day 0). For all groups, S-DNA was photo-conjugated to the hydrogel after sterilization and prior to cell seeding using the previously described method. Three temporal groups were evaluated: OGP-DNA added on Day 0 and removed on Day 7 (Day 0-7, early immobilization); OGP-DNA added on Day 7 with no removal (Day 7-14, delayed immobilization); and OGP-DNA added on Day 0 with no removal (Day 0-14, persistent). OGP-DNA was added on the specified day, in the presence of cells, by removing the cell culture media, adding the complementary OGP-DNA dropwise to the hydrogel surface, and soaking for 5 minutes to allow hybridization to S-DNA. Unbound OGP-DNA was removed via washing with PBS and new cell culture media was added. For the early immobilization group (Day 0-7), OGP-DNA was removed on Day 7 using a similar procedure as addition, except using the cD-DNA for displacement instead of the OGP-DNA. The ability to add and remove cB-DNA in the presence of cells was confirmed using fluorophore labeled-cB-DNA and imaged as previously described. Within each group, equal amounts of S-DNA and OGP-DNA were used, while the cD-DNA concentration that was used was twice the S-DNA/OGP-DNA concentration. The following concentrations were investigated: 0, 0.1, 1, or 10 nm OGP-DNA. On Day 14, alkaline phosphatase (ALP) activity was measured as an early marker for osteogenesis (n=4). Briefly, cells were washed in PBS and lysed using a radio-immunoprecipitation assay (RIPA) lysis and extraction buffer (Thermo Scientific). The cell lysate was incubated with the 1-Step p-nitrophenyl phosphate disodium salt (PNPP) substrate (Thermo Scientific) according to the manufacturer’s protocol. Absorbance at 405 nm was measured and normalized to DNA content using the Quant-iT PicoGreen dsDNA Assay Kit (Invitrogen).

### Statistical Analysis

All data are reported as mean ± standard deviation for at least three independent samples (n=3) unless stated otherwise. For comparing data between experimental groups, a one-way or two-way analysis of variance (ANOVA) followed by Tukey’s post hoc analysis was used to determine statistical significance, where p < 0.05 was considered statistically significant.

Statistical significance is indicated using asterisks. Specifically, * for p < 0.05, ** for p < 0.01, *** for p < 0.001, or **** for p < 0.0001.

## Results and Discussion

### Biomaterial Design and Synthesis

Hyaluronic acid (HA) is a hydrophilic, non-sulfated glycosaminoglycan present throughout the body.^38^ HA is a key component of the extracellular matrix (ECM) and can easily be modified to tailor the resulting material properties as desired.^24,25^ Here, we functionalized HA with norbornene moieties (NorHA) to enable photo-mediated, radical-induced thiol-norbornene click reactions. NorHA functionalization was determined using ^1^H NMR (Figure S1) and was approximately 68%. NorHA was crosslinked using dithiothreitol (DTT) to form a hydrogel for use as a scaffold throughout this study. Hydrogels are three-dimensional, crosslinked polymer networks that can retain a large amount of water and mimic the high water content of native tissues.^39^ NorHA hydrogels were fabricated such that only 20% of the norbornene groups were reacted during the crosslinking step and the remaining norbornene groups were available for reaction with the thiolated DNA after crosslinking. Hydrogels were crosslinked for 5 minutes to ensure complete reaction of the encapsulated crosslinker and prevent further crosslinking with additional UV exposure, which would impact hydrogel mechanical properties. The crosslinking time was selected based on previous research using this hydrogel system, which demonstrated complete reaction within 5 minutes of UV exposure.^40^

To investigate the ability of reversible DNA handles to independently control biomolecule presentation at user-defined time points, four orthogonal sets of single stranded DNA were designed (Table 1). Each set contained the following three oligonucleotides: surface (S)-DNA, complementary biomolecule (cB)-DNA, and complementary displacement (cD)-DNA. The S-DNA contained a thiol functional group to enable thiol-ene photoconjugation to the NorHA hydrogel. The cB-DNA was designed to contain 14 nucleotides complementary to S-DNA to enable cB-DNA immobilization, as well as a toehold region consisting of an additional 5 nucleotides. The cD-DNA was designed to complement the entire cB-DNA sequence, including the toehold region, (19 nucleotides in total) to facilitate cB-DNA removal via toehold-mediated strand displacement. A spacer of 9 nucleotides was included in the S-DNA between the thiol group and the complementary sequence (i.e., hybridization domain) to improve accessibility. The fluorescein-DNA and rhodamine-DNA sets were used to visualize the addition and removal of cB-DNA, including a set of mismatched strands to demonstrate hybridization specificity.

Peptide-DNA conjugates were synthesized using copper-free click chemistry (Figure S2). The peptide-DNA conjugates were purified via reverse-phase high performance liquid chromatography (HPLC) and matrix-assisted laser desorption/ionization time-of-flight (MALDI-TOF) mass spectrometry was used to confirm conjugate identity and purity (Figure S2). Two peptide-DNA conjugates were synthesized for this study: the cell-adhesive peptide RGD (derived from fibronectin)^34^ and the naturally occurring osteogenic growth peptide (OGP).^29,30^ RGD was selected due to its wide use for promoting cell-matrix adhesion on biomaterial scaffolds.^26^ OGP was selected for its ability to enhance osteogenic differentiation of mesenchymal stem cells (MSCs).^31,41^

### Reversible DNA Handles Enable Biomolecule Addition and Removal at User-Defined Timepoints over Multiple Cycles

We evaluated the ability of DNA handles to add and remove biomolecules with complete reversibility and high specificity using rhodamine labeled cB-DNA. First, the S-DNA was photo-conjugated to the NorHA hydrogel using thiol-ene chemistry. The S-DNA served as a permanent handle for the addition and removal of cB-DNA. For addition, a solution of cB-DNA was added onto the hydrogel surface and immobilized via hybridization with the S-DNA on the hydrogel. For removal, a solution of cD-DNA was added onto the hydrogel surface and cB-DNA was removed via toehold-mediated strand displacement. The resulting cB-DNA and cD-DNA duplex was removed via washing and the S-DNA was regenerated for later use. The biomolecule addition and removal scheme is detailed in Figure 1A, with lower case Roman numeral labels indicating points where fluorescence was measured. When S-DNA was present on the hydrogel, the addition of cB-DNA resulted in a high fluorescent signal indicating successful immobilization of cB-DNA onto the hydrogel (i). Next, minimal fluorescent signal was observed following the addition of cD-DNA, indicating successful removal of cB-DNA via toehold-mediated displacement (iv).

Mismatched DNA strands were used to demonstrate the specificity of these interactions (Figure 1B-D). Critically, when cB-DNA was immobilized on the hydrogel, the addition of the mismatched displacement (mD)-DNA did not significantly reduce the fluorescent signal of cB-DNA, which demonstrates successful removal is only possible when the displacement DNA strand is fully complementary (ii). Furthermore, when the mismatched biomolecule (mB)-DNA was added to a hydrogel with S-DNA, the fluorescent signal remained low, which demonstrates successful biomolecule addition requires the biomolecule DNA strand complement S-DNA (iii). When there was no S-DNA on the hydrogel, the addition of cB-DNA did not result in an increase in the fluorescent signal (v). This suggests S-DNA was necessary for successful cB-DNA immobilization and non-specific binding of cB-DNA to the hydrogel did not occur. The fluorescence intensity for each of these experiments is compared in Figure 1E.

Lastly, to evaluate the ability of DNA handles to reversibly add and remove biomolecules, cB-DNA and cD-DNA were sequentially added to the hydrogel over multiple cycles. Fluorescence intensity was measured after each addition of cB-DNA (‘signal ON’) and cD-DNA (‘signal OFF’) to complete five cycles of biomolecule addition and removal. Critically, the fluorescence intensity was relatively consistent for the ‘signal ON’ state, which demonstrates complete reversibility over biomolecule presentation for many cycles. Together, these studies showcase DNA’s unique ability to serve as a highly specific and reversible handle for modulating biomolecule presentation.

### User-Defined, Orthogonal Addition and Removal of Multiple Biomolecules

DNA is advantageous as a reversible handle for multiple biomolecules due to the high specificity of base pairing combined with the vast number of orthogonal oligonucleotide sequences available for use.^42^ We explored the ability of our DNA-based HA hydrogel platform to enable the dynamic presentation of multiple biomolecules using two sets of orthogonal, fluorophore labeled DNA: 1) fluorescein (fluor)-DNA set (numbered set 1) and rhodamine (rho)-DNA set (numbered set 2) (Figure 2). Each set consisted of complementary S-DNA, cB-DNA with the corresponding fluorophore, and cD-DNA as previously described. First, both surface DNA strands (S1-DNA and S2-DNA) were photo-conjugated to the hydrogel. Both complementary biomolecule DNA strands were added to the hydrogel (cB1-DNA with the fluorescein label and cB2-DNA with the rhodamine label) and hybridized to their respective S-DNA. The resulting hydrogel was yellow-orange in fluorescence, which indicates that both fluorophores were present (Figure 2B). Next, the rhodamine signal was selectively removed by adding cD2-DNA (Figure 2C). Critically, the toehold regions between each set were designed to be orthogonal and enable selective removal of the desired biomolecule. The highly selective nature of biomolecule removal was successfully demonstrated by removal of the red fluorescence (cB2-DNA), while retaining the green fluorescence (cB1-DNA). Lastly, the fluorescein signal was added and the rhodamine signal was removed in a single step through simultaneous addition of cD1-DNA and cB2-DNA (Figure 2D). The ability to selectively add and remove multiple biomolecules in a single step showcases the versatility of this approach as a dynamic platform for modulating the presentation of multiple biomolecules. Such a system has the potential to mimic the complex temporal variations in biochemical signals during critical biological processes (e.g., development, healing, and disease progression).

**Figure 2:**
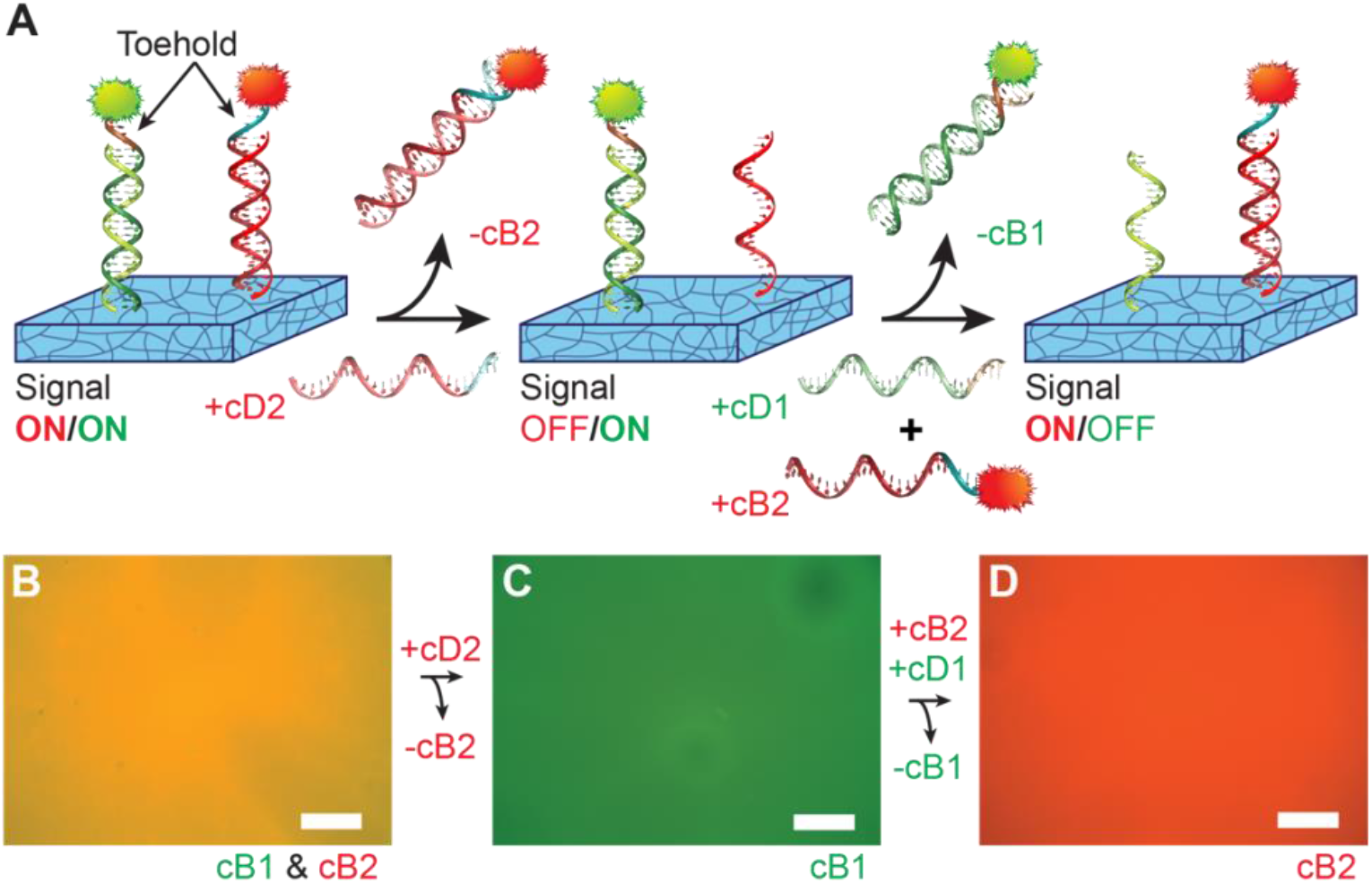
DNA handles enable independent and orthogonal addition and removal of multiple biomolecules at user-defined timepoints. (A) The schematic and corresponding fluorescence image for: (B) two biomolecule signals on the hydrogel (cB1-DNA+S1-DNA in green and cB2-DNA+S2-DNA in red) resulting in a yellow-orange color; (C) addition of cD2-DNA to remove cB2-DNA resulting in a green color; and (D) combined addition of cB2-DNA and cD1-DNA to remove cB1-DNA resulting in a red color (scale = 200 μm).

Next, it was important to verify that biomolecule addition did not impact hydrogel mechanical properties. Biomaterial mechanical properties are known to influence cell behavior, including cell spreading and differentiation.^43–45^ Thus, it is important the hydrogel mechanical properties are not unintentionally altered, which could confound the results. Compression testing was performed to determine the elastic modulus for 1) hydrogels without DNA, 2) S1-DNA and S2-DNA photo-conjugated onto the hydrogel, and 3) S1-DNA+cB1-DNA and S2-DNA+cB2-DNA duplexes immobilized onto the hydrogel. All hydrogel groups had an elastic modulus of approximately 15 kPa and there was no statistical difference between any of the groups (Figure S3). Critically, this result confirms photo-conjugation does not inadvertently add additional crosslinks within the hydrogel, and that the addition of pendent peptide-DNA conjugates does not affect the hydrogel modulus. Other researchers have also performed atomic force microscopy on NorHA hydrogels with and without peptide photoconjugation and did not observe any changes in mechanical properties with peptide addition.^32^

### Temporal Control over RGD Immobilization Modulates Cell Morphology

Cell adhesion allows cells to probe their microenvironment and influences a wide range of cell functions, including proliferation, migration, and differentiation.^46,47^ The role of adhesion in modulating cell behavior has motivated researchers to incorporate cell adhesive ligands into biomaterial scaffolds for a wide range of applications. The most common cell adhesive ligand is RGD, which is derived from fibronectin.^26,34^ Furthermore, cell adhesion, particularly using RGD, has been indicated as a key mediator of osteogenesis.^48,49^ With this in mind, we sought to temporally control RGD immobilization as a tool for modulating cell morphology. First, we confirmed the efficiency of the photoinitiated thiol-ene click reaction between pendent thiol groups and the norbornene groups on the NorHA hydrogel using a thiol quantification assay.

Solutions of the RGD peptide ranging between 0-500 μM were added onto the hydrogel and conjugated to NorHA via exposure to UV light for 60 seconds. After conjugation, the remaining solution was removed and the unreacted RGD concentration was quantified. For all solution concentrations, very little RGD remained after photoconjugation; thus, the tethered RGD concentration was very close to the solution RGD concentration (Figure 3). Thiol-ene reactions are known to be highly efficient, which is one of the advantages of this system.^50,51^

**Figure 3:**
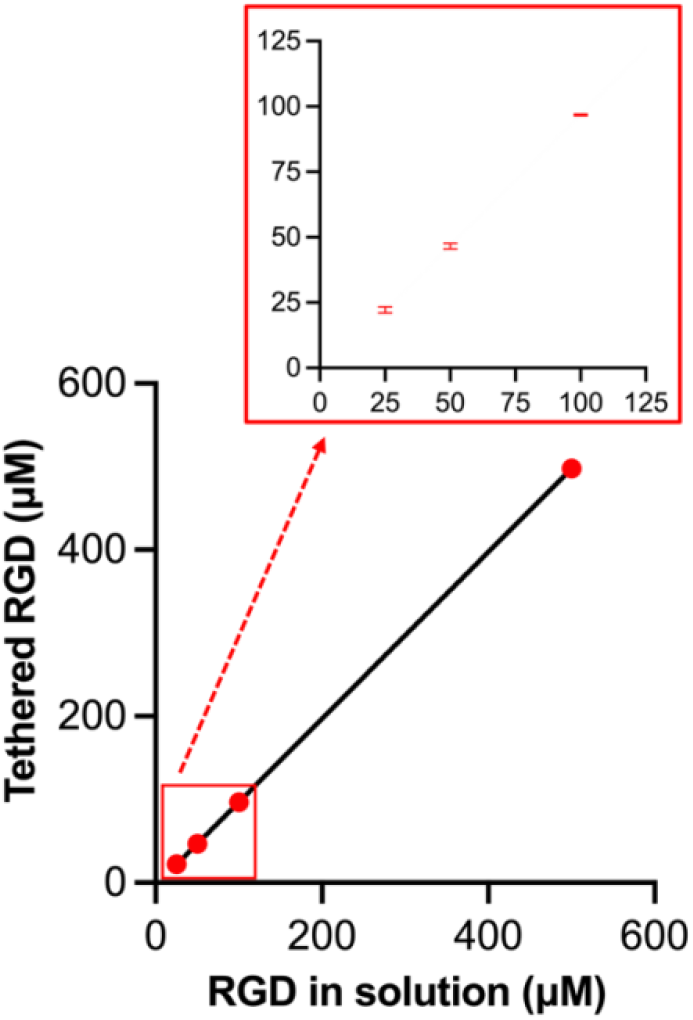
Photoconjugation via norbornene-thiol reaction was highly efficient with complete tethering of the biomolecule to the hydrogel. RGD concentration tethered to hydrogel as a function of initial RGD solution concentration added to the hydrogel, with inset for lower RGD concentrations. Linear regression: y = 1.002x - 3.205, p < 0.0001 (n=3). Error bars represent standard deviations from the mean and are obscured by data points in some cases.

Next, we compared the bioactivity of the RGD-DNA conjugate immobilized onto the hydrogel via DNA handles to the RGD peptide directly photo-conjugated to the hydrogel. The RGD peptide was photo-conjugated onto the hydrogel at a range of concentrations: 0, 100, and 1000 μM. These RGD peptide concentrations were selected based on previous research demonstrating significant cell attachment and spreading within this range on HA biomaterials.^27,28,52^ The RGD-DNA peptide was immobilized onto the hydrogel via hybridization with the complementary S-DNA at 100 μM. This low concentration was chosen for comparison to balance RGD bioactivity with peptide-DNA synthesis constraints. Mesenchymal stem cells (MSCs) were seeded onto hydrogel scaffolds with or without RGD and cell morphology was evaluated after 3 days of in vitro cell culture. Cells were stained with phalloidin to visualize the cytoskeleton and calculate cell area and DAPI to visualize the cell nucleus and calculate cell number (Figure 4). As expected, a larger cell area was observed on hydrogels with a higher RGD concentration (1000 > 100 > 0 μM). Also, significantly more cells were attached onto hydrogels with the highest RGD concentration (1000 μM) compared to all other groups. No statistical difference in cell area or number was noted between the RGD peptide tethered directly and the RGD-DNA conjugate immobilized onto the hydrogel. Although the RGD-DNA conjugate promoted a slightly higher cell area compared to the RGD peptide, this difference was not significant. The RGD-DNA conjugate (100 μM) promoted a statistically significant higher cell area compared to no peptide and was chosen for the temporal variation studies. Critically, HA does not innately support integrin-mediated cell adhesion; thus, we can precisely modulate cell adhesion on the hydrogel scaffolds via the inclusion of cell adhesive ligands. As such, limited cell attachment and spreading was expected in the absence of RGD.

**Figure 4:**
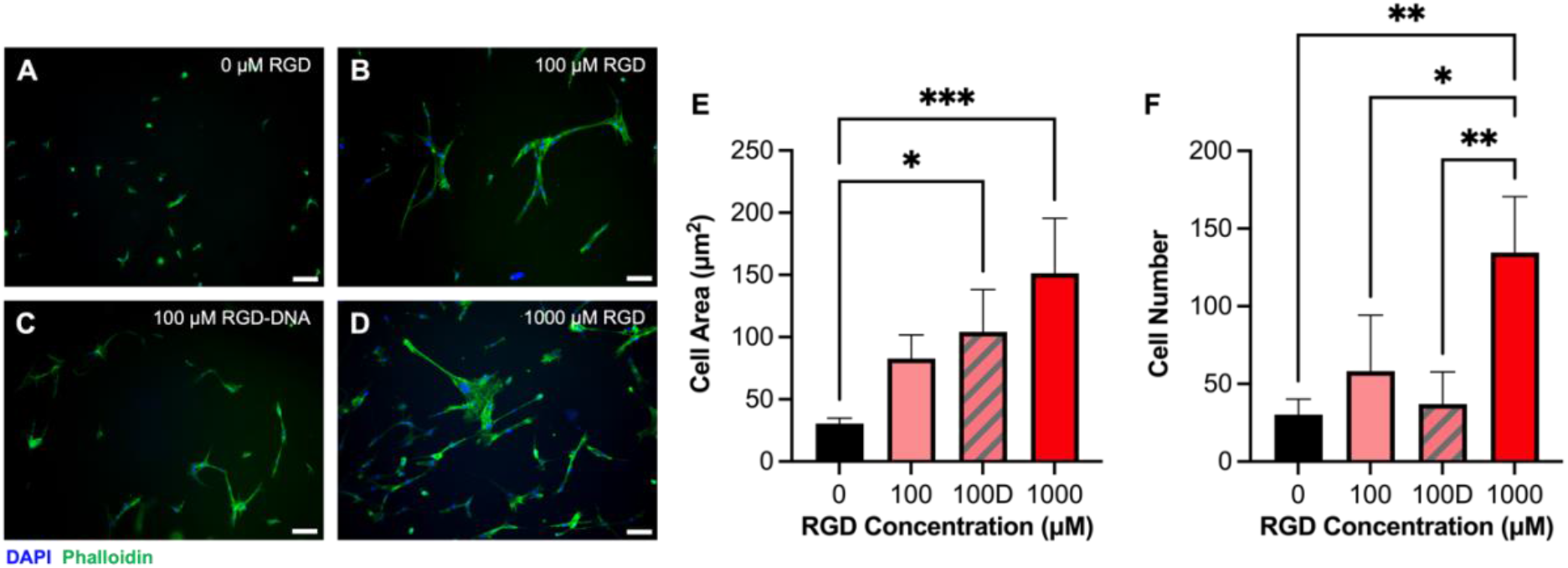
The cell adhesive peptide, RGD, promoted cell spreading and was not impacted by the inclusion of DNA as a reversible handle. Mesenchymal stem cells (MSCs) were seeded onto hydrogels containing (A) 0 μM RGD, (B) 100 μM RGD, (C) 100 μM RGD-DNA, or (D) 1000 μM RGD and stained with DAPI (blue) and phalloidin (green) to visualize the cell nucleus and cytoskeleton, respectively (scale = 200 μM). Cell (E) area and (F) number as a function of RGD concentration (n=3). Asterisks indicate statistical significance as noted in the statistical analysis methods. Error bars represent standard deviations from the mean.

To demonstrate the utility of this platform to temporally modulate cell morphology, we cultured MSCs on hydrogels containing the immobilized RGD-DNA conjugate, removed RGD-DNA on day 3 of cell culture through addition of cD-DNA, and assessed cell morphology on day 7 of cell culture. Temporal variation was compared to the persistent presentation of the RGD-DNA conjugate (i.e., RGD-DNA present for all 7 days of the cell culture study). The ability to add and remove cB-DNA in the presence of cells was confirmed using fluorophore labeled cB-DNA (Figure S4). Scaffolds were stained with phalloidin (cytoskeleton) to calculate cell area and aspect ratio, as well as DAPI (cell nucleus) to calculate cell number (Figure 5). Cells were moderately spread with a slightly elongated aspect ratio on hydrogels with persistent RGD-DNA presentation after 7 days of cell culture. When the RGD-DNA conjugate was removed on day 3, cells contracted with a significant decrease in cell area and aspect ratio by day 7 of cell culture.

**Figure 5:**
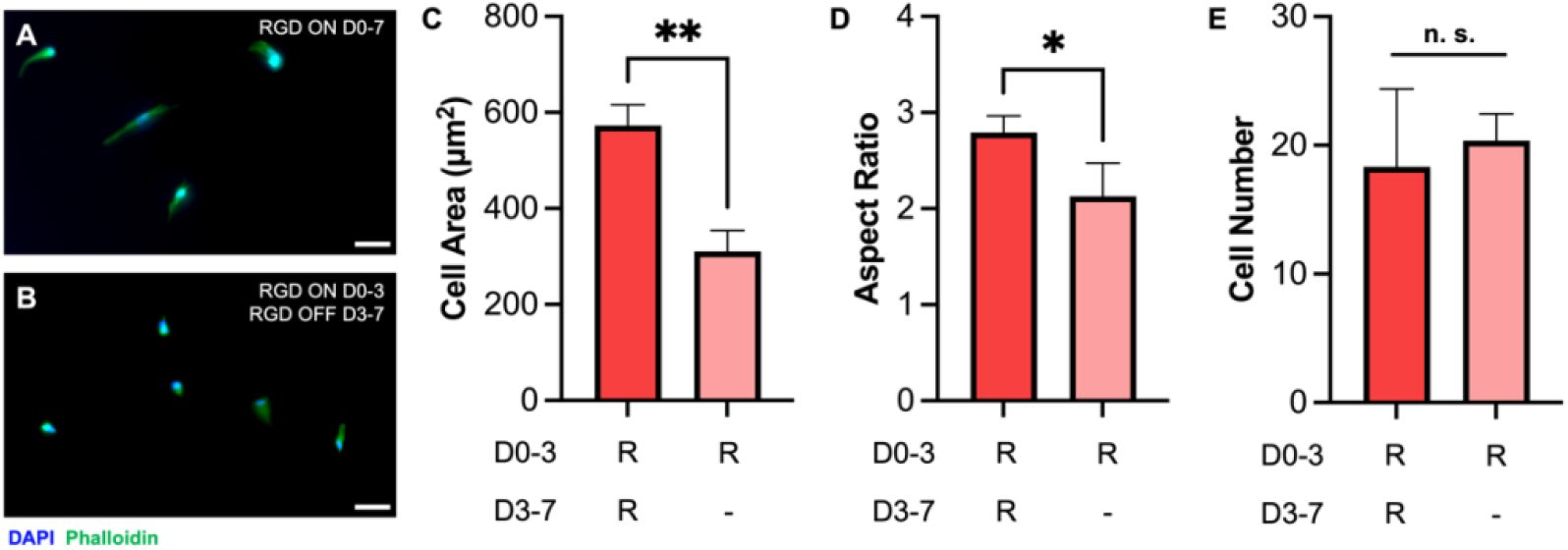
User-controlled addition and removal of the cell adhesive peptide, RGD, enables temporal control over cell spreading. Mesenchymal stem cells (MSCs) were seeded onto hydrogels with (A) RGD on from day 0-7 or (B) RGD on from day 0-3 and off from day 3-7. All hydrogels were stained on day 7 with DAPI (blue) and phalloidin (green) to visualize the cell nucleus and cytoskeleton, respectively (scale = 200 μM). Cell (C) area, (D) aspect ratio, and (E) number were quantified for both experimental groups (n=3). Asterisks indicate statistical significance as noted in the statistical analysis methods and n. s. indicates no statistical significance. Error bars represent standard deviations from the mean.

At this low RGD-DNA concentration (100 μM), no statistical differences were noted in cell number between groups, which was similar to the trend observed in Figure 4F. This behavior corroborates previous research that has observed a significant reduction in cell area following the removal of a cell adhesive ligand.^20,52^

#### Delayed Presentation of OGP Increases Osteogenesis

Osteogenic growth peptide (OGP) is a naturally occurring growth factor present in human blood serum. OGP is composed of 14 amino acid motifs which are identical to the C-terminus of histone H4’s amino acid sequence.^29,30^ OGP is proteolytically degradable, resulting in the C-terminal pentapeptide termed OGP(10-14). OGP(10-14) is considered the bioactive domain, since this sequence is known to activate the mitogen-activated protein (MAP) kinase signaling pathway.^53^ Natively, OGP is present in a variety of forms including: protein bound OGP, soluble OGP, and proteolytically cleaved soluble OGP(10-14).^30^ Significant research has shown that OGP plays a critical role during bone repair through the stimulation of proliferation, osteoblast differentiation, and alkaline phosphatase (ALP) activity.^30,41,53,54^ Thus, OGP is a promising candidate to promote osteogenesis, where immobilization of OGP using DNA handles mimics the naturally occurring protein bound form of OGP.

To evaluate the effect of OGP concentration on osteogenesis, MSCs were seeded onto hydrogels with varying concentrations of either OGP photo-conjugated directly to the hydrogel or OGP-DNA immobilized onto the hydrogel using DNA handles. The OGP concentration ranged between 0.1 to 10 nM and was compared to hydrogels without OGP. This concentration range was selected based on previous research which has reported 1 nM was the ideal OGP concentration for enhancing ALP activity in human primary osteoblast cells.^55^ Cells were exposed to persistent OGP presentation and ALP activity was measured after 14 days of in vitro cell culture (Figure 6A). All hydrogels with OGP, regardless of concentration or immobilization method, promoted statistically significant ALP activity compared to hydrogels without OGP. Minimal differences in ALP activity were noted between OGP concentrations and immobilization method. That being said, the ALP activity for OGP photo-conjugated directly to the hydrogel was highest at 1 nM OGP and significantly decreased at higher OGP concentrations (10 nM). This behavior is similar to the trends reported previously.^55^ The abrogation of this effect when presenting OGP using DNA handles, may indicate the dynamic presentation of OGP via DNA hybridization is beneficial for osteogenesis.

**Figure 6:**
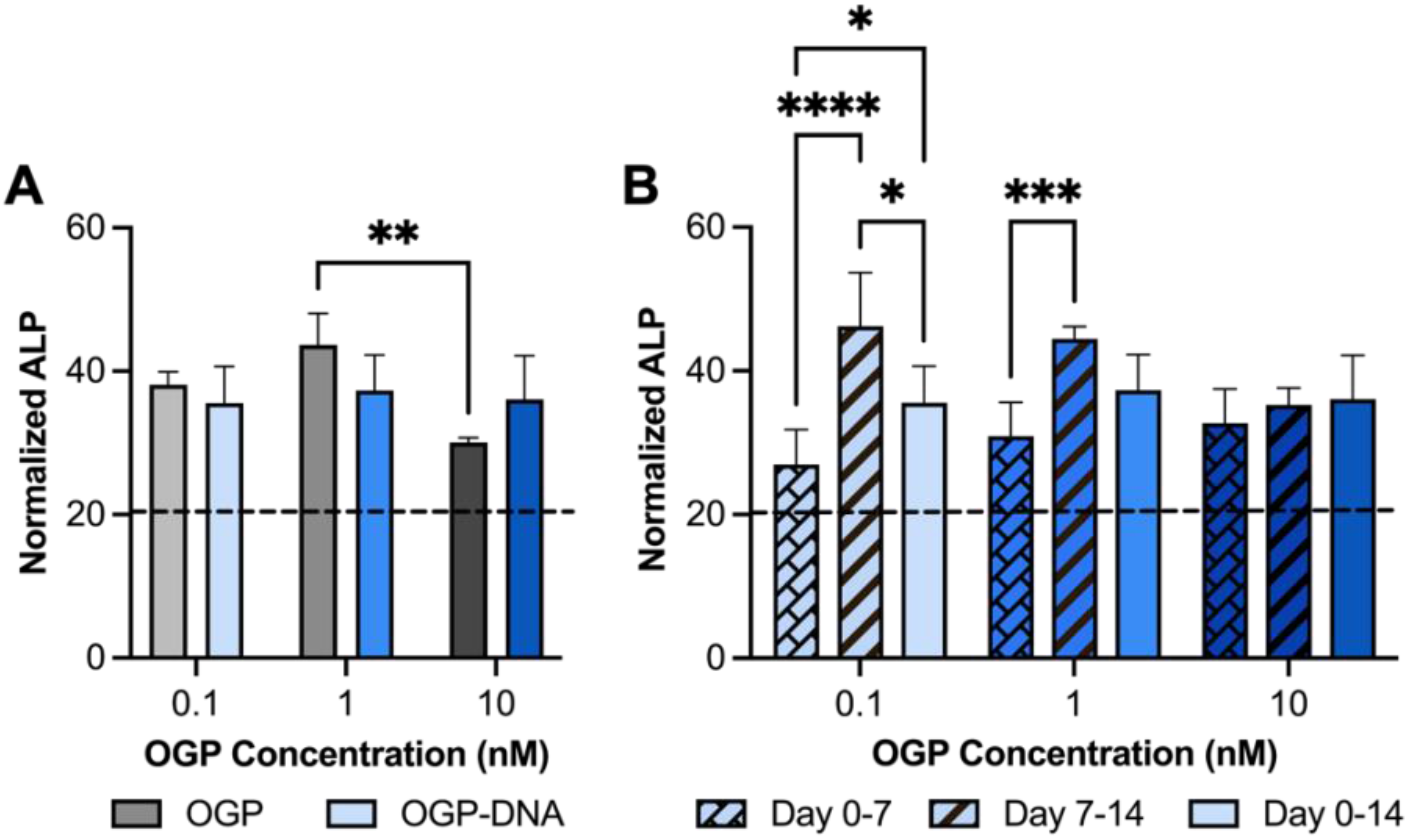
Delayed presentation of osteogenic growth peptide (OGP) increases alkaline phosphatase (ALP) expression. (A) ALP expression on day 14 following sustained presentation of OGP using either the OGP peptide or the OGP-DNA conjugate and as a function of OGP concentration. (B) ALP expression on day 14 as a function of OGP-DNA concentration and temporal presentation. Three OGP-DNA temporal presentation groups were compared: OGP on from day 0-7 and off from day 7-14; OGP off from day 0-7 and on from day 7-14; and OGP on from day 0-14. Asterisks indicate statistical significance as noted in the statistical analysis methods (n=4). For visual simplicity, statistical significance is only shown in (B) for comparisons within each concentration. Dashed lines indicate average ALP expression on day 14 for hydrogels with no OGP. Error bars represent standard deviations from the mean.

To evaluate the temporal role of OGP during osteogenesis, we used DNA handles to add and remove OGP-DNA at user-defined times. Three temporal groups were compared: early immobilization (OGP on from day 0-7 and off from day 7-14); delayed immobilization (OGP off from day 0-7 and on from day 7-14); and persistent immobilization (OGP on day 0-14). Cells were seeded onto hydrogels with or without OGP and ALP activity was measured for all groups after 14 days of in vitro cell culture (Figure 6B). Interestingly, ALP activity was significantly higher when OGP presentation was delayed compared to early or persistent immobilization.

This trend was only observed for lower OGP concentrations tested (0.1 and 1 nM OGP) and was not present in the highest OGP concentration tested (10 nM OGP). The lack of a relationship between OGP temporal presentation and ALP activity at higher OGP concentrations may be related to the negative regulation observed at supraoptimal OGP doses (>1 nM).^55^ All ALP activity was normalized to DNA content. DNA content for all experimental groups is summarized in Figure S6. No consistent trends in DNA content were observed as a function of OGP concentration, immobilization method, or temporal presentation. Together, these results suggest selecting the optimal concentration and temporal presentation of OGP is critical in promoting OGP-induced osteogenesis for bone repair applications.

## Conclusions

Due to the highly programmable nature of DNA, it is an ideal linker for the tethering of biomolecules onto biomaterial scaffolds. We combined reversible DNA handles with norbornene-modified HA, where thiol-ene chemistry was used to orthogonally photo-conjugate one or more surface DNA strand(s). Next, we were able to immobilize bioactive peptides of interest by adding a complementary biomolecule-DNA conjugate that hybridized with the surface DNA strand. The biomolecule-DNA conjugate was removed using toehold-mediated displacement through the addition of a complementary displacement-DNA stand. Using this approach, we demonstrated orthogonal control over the addition and removal of multiple biomolecules over many cycles. We used this platform to investigate the relationship between temporal peptide presentation and cell behavior for two peptides of interest: the cell adhesion peptide RGD and osteogenic growth peptide (OGP). First, we observed that persistent RGD presentation is important for promoting cell spreading, with cells contracting when RGD was removed. Second, we observed delayed OGP presentation increased alkaline phosphatase activity compared to other temporal experimental groups. Together, these results highlight the importance of optimizing the temporal presentation of biochemical signals during tissue regeneration. Critically, these results also demonstrate we were able to modulate cell behavior through the addition or removal of biochemical signals during long-term in vitro cell culture (up to 14 days). This DNA-based hydrogel platform can be easily adapted to temporally modulate a variety of peptides and other biomolecules of interest. Further, the thiol-ene photoconjugation step enables orthogonal spatial patterning of the surface DNA strand(s),^32^ which could be employed to precisely control biomolecule presentation in space and time. Thus, this modular platform provides a unique approach to tease apart the spatiotemporal role of multiple biomolecules during development, regeneration, and disease progression.

## Acknowledgements

The authors kindly acknowledge funding from the National Institutes of Health (NIH): National Institute of Arthritis and Skin Diseases (NIAMS) under award R21AR074069 (J.L.H and N.S.) and National Institute of General Medical Sciences (NIGMS) under award DP2GM132931 (N.S.). Additional funding was provided from the National Science Foundation (NSF): DMR-BMAT under CAREER award 1753387 (N.S.). The content in this manuscript is solely the responsibility of the authors and does not necessarily represent the official views of NIH or NSF. Instrumentation support was provided in part by ASU’s Advanced Light Microscopy Core Facility, Regenerative Medicine Bioimaging Core Facility, and the Magnetic Resonance Research Center.

