## Supplemental Information for "Using dynamic biomaterials to study the temporal role of osteogenic growth peptide during osteogenesis"

### Supplementary Information

Fallon M. Fumasi[a], Tara MacCulloch[b,c], Julio Bernal-Chanchavac[b,c], Nicholas Stephanopoulos[b,c], and Julianne L. Holloway[a,b]

[a]Chemical Engineering; School for Engineering of Matter, Transport and Energy; Arizona State University; Tempe, Arizona

[b]Biodesign Center for Molecular Design and Biomimetics; Arizona State University; Tempe, Arizona

[c]School of Molecular Sciences; Arizona State University; Tempe, Arizona

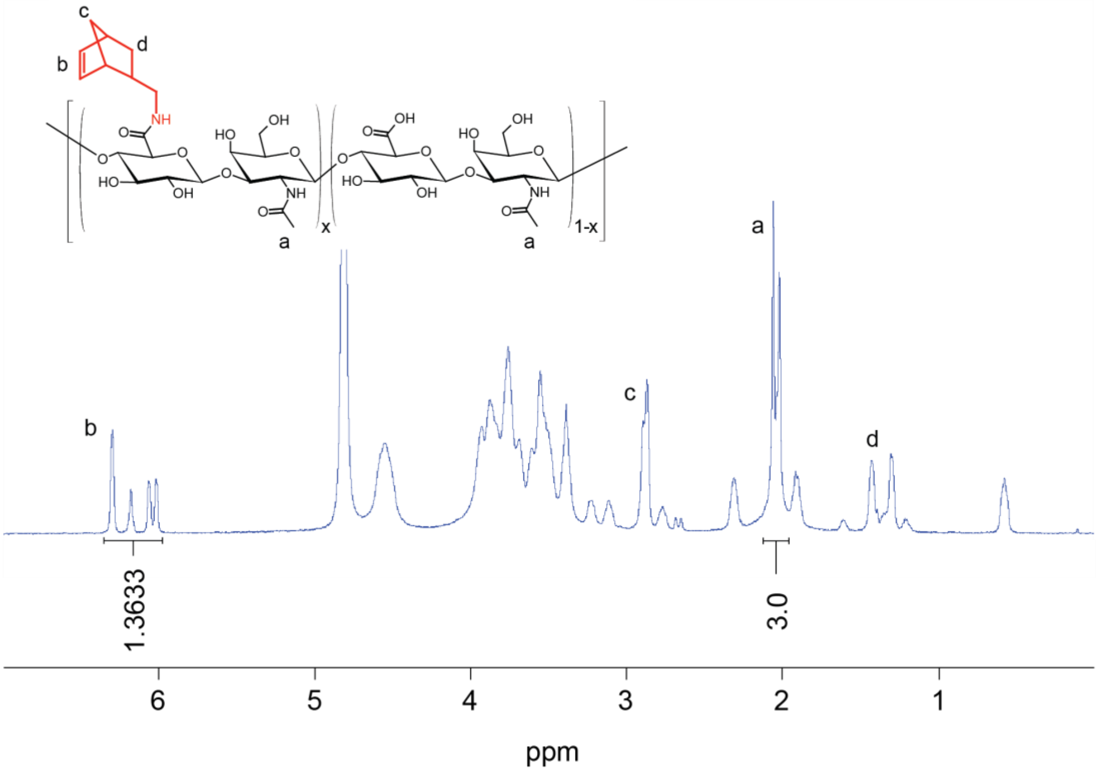


**Figure S1: NorHA chemical structure and functionalization was quantified using $^{1}H$ NMR.** $^{1}H$ NMR spectrum for NorHA, where peaks associated with the norbornene group between 6.0 and 6.3 ppm (2H, C=C, b) and the peak associated with the methyl group on the hyaluronic acid backbone at 2.06 ppm (3H, CH3, a) were used to calculate functionalization. Additional peaks associated with the norbornene group are also indicated at 2.87 ppm (CH2, c) and between 1.3-1.43 ppm (CH2, d). In this spectrum, HA was 68% functionalized with norbornene groups.

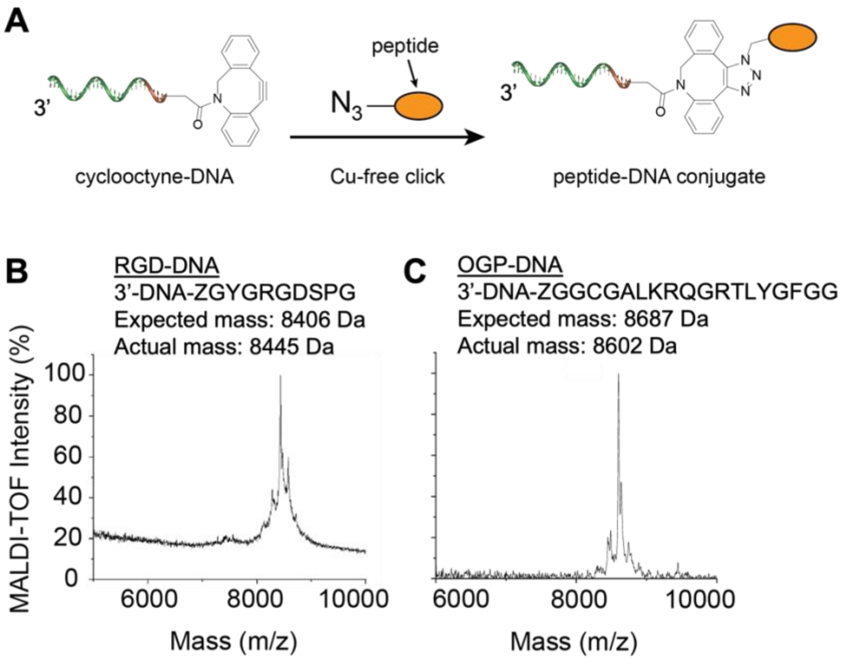


**Figure S2: MALDI-TOF was used to confirm peptide-DNA conjugate formation.** (A) Peptide-DNA conjugates were synthesized by reacting an azidolysine on the peptide with a dibenzocyclooctyne on the DNA via copper-free click chemistry. MALDI-TOF mass spectrometry was used to determine conjugate mass and purity for (B) RGD-DNA and (C) OGP-DNA. Possible discrepancies between the expected and actual mass may be due to trace amounts of salts (e.g., sodium, potassium, etc.), calibration errors, or differences in the DNA mass ordered and the actual DNA mass.

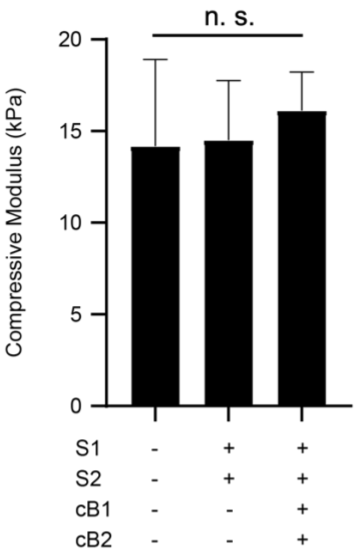


**Figure S3: The addition of peptide-DNA conjugates onto the NorHA hydrogel did not alter hydrogel mechanical properties.** Hydrogel compressive modulus was measured for: 1) hydrogels without DNA, 2) S1-DNA and S2-DNA photo-conjugated onto the hydrogel, and 3) S1-DNA+cB1-DNA and S2-DNA+cB2-DNA duplexes immobilized onto the hydrogel. No statistical differences between experimental groups were observed ($p > 0.05$) (n=6). Error bars represent standard deviations from the mean.

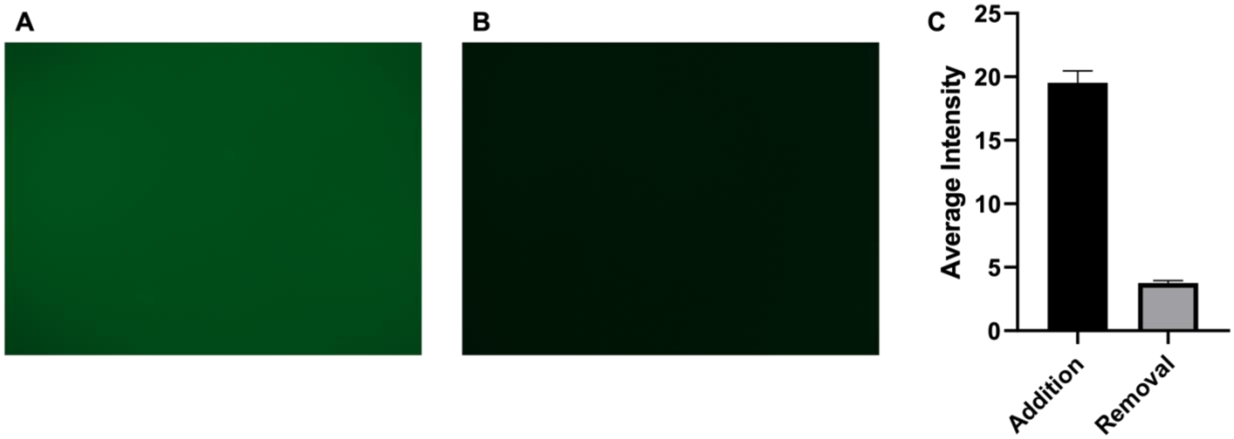


**Figure S4: Peptide-DNA conjugates were successfully added and removed in the presence of cells.** (A) Fluorescein labeled cB-DNA was added onto the hydrogel in the presence of cells and (B) removed from the hydrogel in the presence of cells via addition of cD-DNA. (C) Fluorescence intensity was visualized and quantified after addition and removal (n=5). Error bars represent standard deviations from the mean.

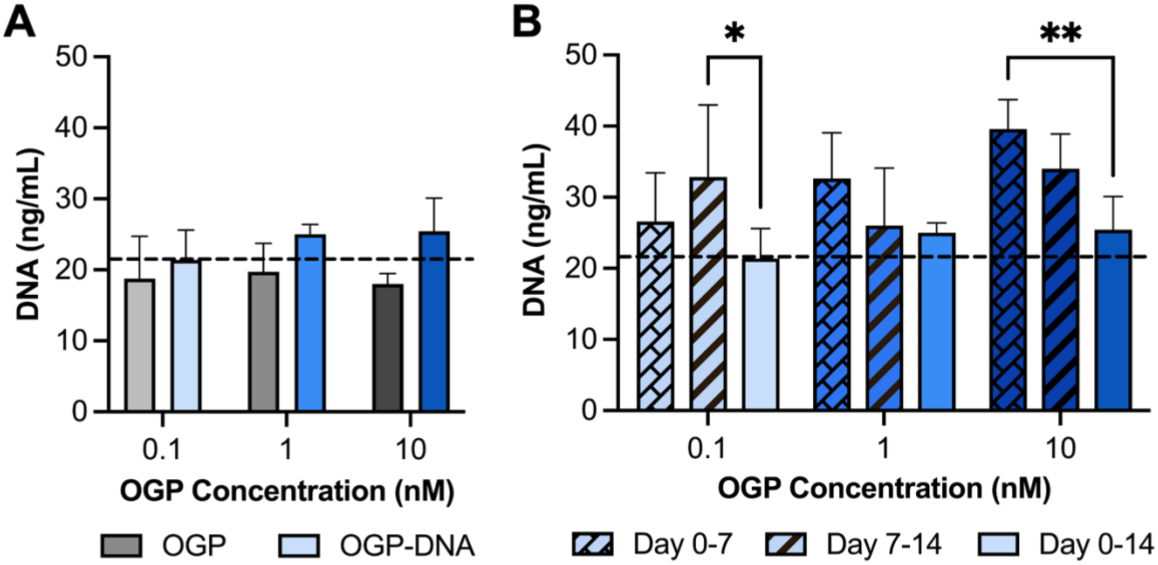


**Figure S5: No consistent trends in the number of adhered cells, as indirectly evaluated using DNA quantification, were observed as a function of OGP concentration or temporal presentation.** (A) DNA quantification on day 14 following sustained presentation of OGP using either the OGP peptide or the OGP-DNA conjugate and as a function of OGP concentration. (B) DNA quantification on day 14 as a function of OGP-DNA concentration and temporal presentation. Three OGP-DNA temporal presentation groups were compared: OGP on from day 0-7 and off from day 7-14; OGP off from day 0-7 and on from day 7-14; and OGP on from day 0-14. Asterisks indicate statistical significance as noted in the statistical analysis methods (n=4). For visual simplicity, statistical significance is only shown in (B) for comparisons within each concentration. Dashed lines indicate average DNA quantification on day 14 for hydrogels with no OGP. Error bars represent standard deviations from the mean.
